# Deciphering landscape-scale plant cover and biodiversity from soil eDNA

**DOI:** 10.1101/2025.01.14.632973

**Authors:** Tim Goodall, Robert I. Griffiths, Hyun S. Gweon, Lisa Norton, Susheel Bhanu Busi, Daniel S. Read

## Abstract

Biodiversity surveys are critical for detecting environmental change; however, undertaking them at scale and capturing all available diversity through observation is challenging and costly. This study evaluated the potential of soil-extracted eDNA to describe plant communities and compared these findings to traditional, observation-based, field surveys. We analysed 789 soil samples using high-throughput amplicon sequencing and compared DNA-based diversity metrics, indicator taxa, predicted vegetation class, and plant cover in a comparison with co-located field survey data. The results indicated that taxonomically aggregated (genus) eDNA-derived data, while showing slightly reduced Shannon’s diversity scores, yielded remarkably similar overall richness and composition estimates. However, the DNA indicator taxa and predictive power for vegetation community classification were also lower overall than those recorded by the field survey. However, in many cases plant cover could be inferred from amplicon abundance data with some accuracy despite widely differing scales of sampling – 0.25 g crumb of soil versus a 1 m^2^ quadrat. Overall, results from eDNA demonstrated lower sensitivity but were broadly in accordance with traditional surveys, with our findings revealing comparable taxonomic resolution at the genus level. We demonstrate the potential and limitations of a simple molecular method to inform landscape-scale plant biodiversity surveys, a vital tool in the monitoring of land use and environmental change.

## Introduction

In the face of accelerating climate change, landscape-scale monitoring of biodiversity is essential for detecting shifts in species composition, ecosystem function, and habitat health^1^, which can have profound implications for ecosystem resilience and the services upon which human societies rely. Landscape scale temporal biodiversity inventories, when combined with comprehensive environmental metadata, allow the observation of ecosystem trends and are vital tools for understanding and estimating habitat change, informing models, and enabling the forecast of future change^2^. Such models can be used to estimate the response of ecosystems and their services to anthropogenic impacts, the effects of climatic change, or even government policy^3^. Arguably, because of the increasing rate of change and our dependence on ecosystem services, monitoring the natural environment through biodiversity assessment has never been as important, yet monitoring is often patchy or piecemeal in coverage and is dependent upon resource availability and legislative drivers^4^.

Traditional approaches to landscape-scale biodiversity surveys require an expansive group of field ecologists to visit hundreds or thousands of locations and to perform consistent and accurate measurements of plant species. Because these surveys only offer a snapshot of visible plant growth, dormant, ephemeral, or cryptic plants may not be observed^5^. Resource availability, both human and financial, is a limiting determinant of survey scale and sampling intensity. New methods to maximise efficiency, facilitate greater sampling depth, and increase the scale of surveys are desirable. At large scale, methods such as drone^6^, aircraft or satellite remote sensing are useful tools^7^, however their resolution is limited. Environmental DNA (eDNA) monitoring offers the potential to simplify the field effort required and enable the processing of vast numbers of samples with high taxonomic resolution^8^.

eDNA analysis has provided insight into many otherwise difficult-to-monitor environments and assists in estimating biodiversity and distribution of both micro-and, more recently, macro-organisms^8,9^. The accuracy of eDNA for plant community analysis from soil is relatively novel and untested at scale. Fahner et al.^10^ assessed a suite of plant taxonomic markers in 35 forest soils, upon which we built herein by applying the best-performing taxonomic markers to examine the ability of soil eDNA to represent local plant communities from different soils across a national landscape. In this study, we extracted eDNA from 789 soil samples collected as part of the UK Centre for Ecology and Hydrology (UKCEH) Countryside Survey of 2007^11^, where each sampling location was simultaneously subjected to vegetative studies by trained plant ecologists. We assessed how data derived from a high-throughput molecular approach and classical field survey methods compared to each other, and explored the merits and limitations of this eDNA approach within the context of a national survey. Specifically, we compared the key indicators of Aggregate Vegetation Classification (AVC) types, the predictive ability of the data to ascribe AVC class using machine learning and examine the potential for molecularly derived abundance data to describe plant cover. Our findings highlight the merits of each approach and, importantly, inform the potential for molecularly derived methods, specifically amplicon-based soil-eDNA, to describe plant cover and overall biodiversity within these ecosystems.

## Methodology

### Vegetation survey

Surveyors undertook vegetation surveys as part of the 2007 UK countryside survey following published guidelines^12^ (Figure 1). For the purposes of this study, we used the 1m^2^ plant species recordings (nest 0). The surveys were conducted at a minimum of one meter and a maximum of two and a half meters distance from the location of the soil sample.

**Figure 1.**
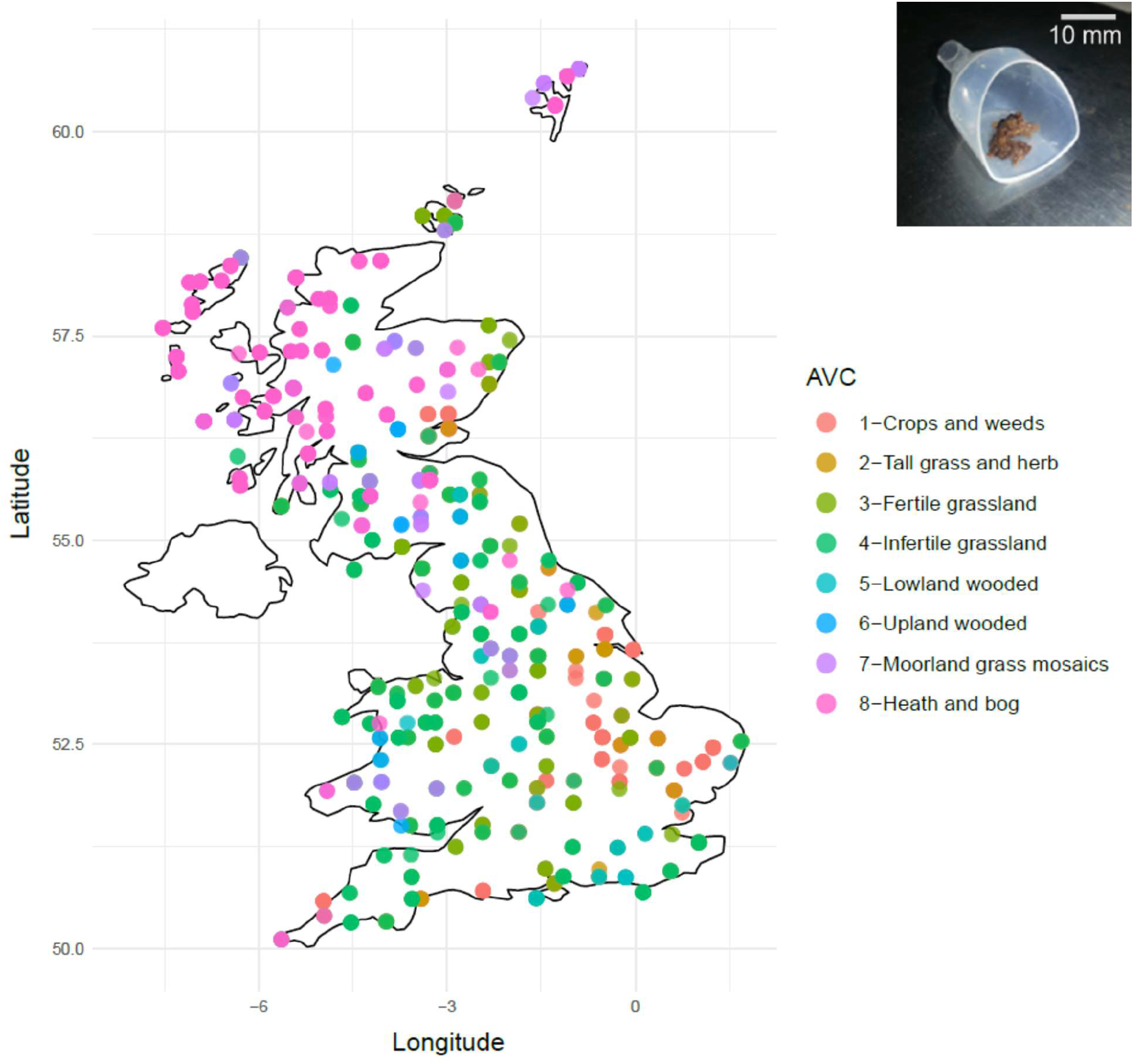
Distribution of survey sites within England, Scotland, and Wales; coloured by AVC classification. For data protection site co-ordinates are rounded to the nearest 10 km. **(inset)** 0.25 g of homogenized soil prior to DNA extraction.

### Soil collection

Soil sampling and vegetation surveys were conducted simultaneously. A five-centimetre diameter soil core was collected from the top 15 cm at each sample site. The cores were transferred to the laboratory and subjected to multiple analyses^13^. Sub-samples of sieved, homogenised, field moist soils were archived at -20 °C for later DNA extraction and plant ITS2 amplicon sequencing.

### Vegetation classification

Vegetation communities are closely aligned with habitat definitions and assessments of habitat health. An Aggregate Vegetation Classification (AVC) was applied to each sample site based on the surveyor’s plant species cover estimates. AVCs are determined as per Bunce et al.^14^; briefly, plant survey data is lumped and subjected to multivariate analysis using DECORANA and ordinated. Clustering of the sample within one of eight groups forms the basis for the classification, where the eight AVC classes are defined as: 1 ‘Crops and weeds’, 2 ‘Tall grass and herb’, 3 ‘Fertile grassland’, 4 ‘Infertile grassland’, 5 ‘Lowland wooded’, 6 ‘Upland wooded’, 7 ‘Moorland grass mosaics’, and 8 ‘Heath and bog’.

### Molecular analyses of plant ITS2

DNA was extracted from 0.25 g of the archived soil (Figure 1 inset) using a Powersoil DNA Isolation Kit (Qiagen Ltd.) according to the manufacturer’s instructions. Amplicons were generated using a 2-step amplification approach, with Illumina Nextera tagged ITS2 region primers, forward primer ITS2-S2 ATGCGATACTTGGTGTGAAT and reverse primer ITS4 TCCTCCGCTTATTGATATGC^10^, each modified at the 5’ end with the addition of Illumina pre-adapter and Nextera sequencing primer sequences. Amplicons were generated using high-fidelity DNA polymerase (Q5 Taq; New England Biolabs). After initial denaturation at 95 °C for 2 min, the PCR conditions were as follows: denaturation at 95 °C for 15 s, annealing at 55 °C, annealing for 30 s with extension at 72 °C for 30 s, repeated for 35 cycles. A final extension step of 10 min at 72 °C was performed. PCR products were cleaned using a Zymo ZR-96 DNA Clean-up Kit (Zymo Research, US) following the manufacturer’s instructions. MiSeq adapters and 8nt dual-indexing barcode sequences were added during the second PCR amplification step. After an initial denaturation at 95 °C for 2 min, the PCR conditions were as follows: denaturation at 95 °C for 15 s, annealing at 55 °C, annealing for 30 s with extension at 72 °C for 30 s, repeated for eight cycles with a final extension of 10 min at 72 °C. Amplicon sizes were determined using an Agilent 2200 TapeStation system. Libraries were normalised using the SequalPrep Normalisation Plate Kit (Thermo Fisher Scientific), quantified using the Qubit dsDNA HS kit (Thermo Fisher Scientific), and pooled. The pooled library was diluted to 400pM after denaturation and neutralisation. Denaturation was achieved with 0.2N NaOH for 5 min, followed by neutralisation with 0.2N HCl. The library was then diluted to a load concentration of 12 pM with HT1 Buffer and a 10% denatured PhiX control library. The final denaturation was performed by heating to 96°C for 2 min, followed by cooling in crushed ice. Sequencing was performed on an Illumina MiSeq using V3 600 cycle reagents. The 789 samples were randomly split over three flow cells, with each generating in excess of 17,000,000 reads passing filter.

### Normality and correlation analysis

Data derived from either survey method were determined by the Shapiro test to be non-normally distributed; therefore, non-parametric Spearman’s correlations were used to assess the relationships between the survey methods. Tests were conducted to compare (i) diversity measures of genera, (ii) richness of the genera recorded, and (iii) AVC indicator genera derived from each survey method. The number of sites examined totalled 789 of which 126 were of AVC-1 ‘Crops and weeds’, 35 of AVC-2 ‘Tall grass and herb’, 173 of AVC-3 ‘Fertile grassland’, 190 of AVC-4 ‘Infertile grassland’, 19 of AVC-5 ‘Lowland wooded’, 28 of AVC-6 ‘Upland wooded’, 69 of AVC-7 ‘Moorland grass mosaics’, and 149 of AVC-8 ‘Heath and bog’.

### Molecular Bioinformatics

Illumina demultiplexed sequences were processed using HONEYPI (https://github.com/hsgweon/honeypi) to generate Actual Sequence Variant (ASV) tables and sequence taxonomies. After passing through HONEYPI taxonomies and ASV tables were merged by ASV sequence.

### Analysis

After quality filtering, matching the outputs of the molecular analysis to field surveys permitted a comparison of methodologies at 789 sites spanning eight AVC classifications. The data were processed at two levels.

Firstly, at genus level (to enable taxonomic comparisons), all non-flowering plant data were removed from both the ASV and survey tables, thus bryophytes, algae, gymnosperms, bare ground, leaf litter and rock were excluded. Counts of species were collapsed to genus level to avoid issues with taxonomic disagreements and ambiguous survey recordings, for example, the misappropriation of sequence reads for Oil-seed Rape (Brassica napus) as both, or either parent line (Brassica rapa or oleracea). Proportional abundances were then calculated for each sample’s molecular data using decostand (R package vegan). Each sample’s rare genera (<5% abundance) were removed before subsequent analysis. To assess the similarities in taxonomic observations between molecular and survey data, each site’s AVC classification was used to determine indicator genera (R package labdsv), Shannon’s diversity and the richness of genera recorded by each survey method (R package vegan). Shapiro tests were performed to determine the normality of the responses. Statistical comparison of eDNA abundance and 1 m2 plant cover survey methods: Spearman’s Rho statistic was calculated to estimate a rank-based measure of association between the survey methods (base R), specifically, the correlation between the abundance of co-recorded genera at each site within each AVC, as well as the correlation of genus level diversity measures at each site and within each AVC.

Secondly, the species level data were used to compare the predictive ability of the eDNA abundance and 1 m^2^ plant cover survey in ascribing the sample site’s AVC classification using machine learning (R package xgboost^15^), with a 4:1 training to testing split, using settings: method = “xgbTree”, tuneGrid = expand.grid(nrounds = c(50, 100), max_depth = c(2, 4, 6), eta = c(0.1, 0.3), gamma = c(0, 1), colsample_bytree = c(0.7), min_child_weight = c(1), subsample = c(0.8), with a 5-fold cross validation check.

## Results

### Measured genus level diversity and richness and correlation between methods

The community-level relative abundance data for flowering plants from each site, collapsed to the genus level, were used to calculate Shannon diversity scores per survey (Figure 2). Score averages were calculated per AVC and the results were, for molecular and 1 m^2^ surveys: Crops and weeds: 0.57 (SD 0.48) and 0.16 (SD 0.35), Tall grass and herb: 0.48 (SD 0.37) and 0.69 (SD 0.58), Fertile grassland: 0.65 (SD 0.43) and 0.8 (SD 0.50), Infertile grassland: 0.85 (SD 0.49) and 1.32 (SD 0.49), Lowland wooded: 0.56 (SD 0.44) and 0.78 (SD 0.46), Upland wooded: 0.35 (SD 0.39) and 0.73 (SD 0.52), Moorland grass mosaics: 0.64 (SD 0.43) and 1.22 (SD 0.53), Heath and bog: 0.38 (SD 0.35) and 1.93 (SD 0.43). Across all samples, the average Shannon’s diversity scores were 0.61 (SD 0.47) and 0.89 (SD 0.61) for molecular and 1 m2 surveys, respectively.

**Figure 2.**
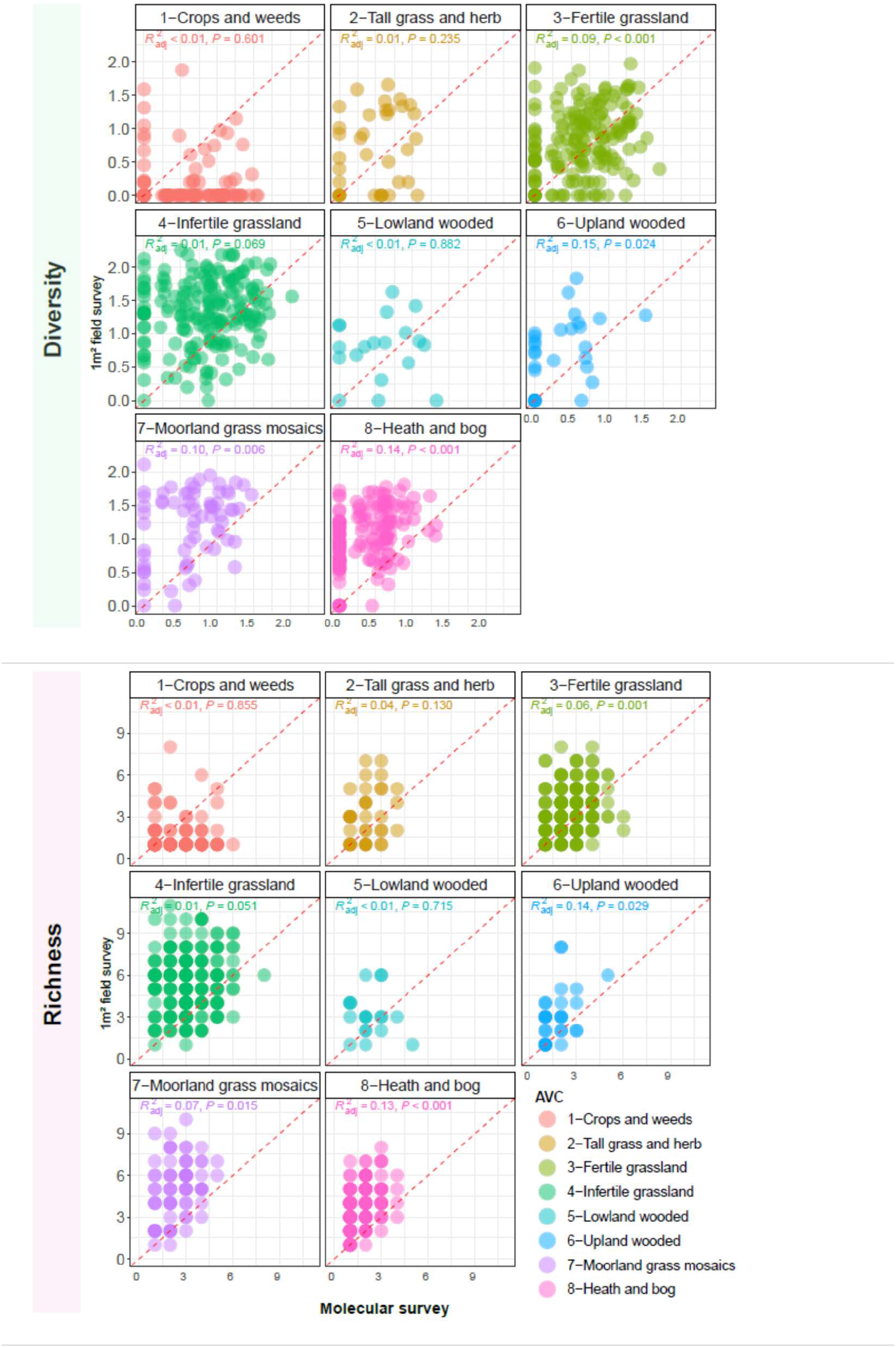
Scatter plots, faceted by AVC, showing each site’s genus level diversity score and richness of genera; 1m^2^ plant cover (X-axis) and molecular surveys (Y-axis) with linear regression values and dotted 1:1 line.

Aside from AVC-1 ‘Crops and weeds’, the molecular survey recorded, on average, lower Shannon’s diversity scores. Spearman’s rank correlation coefficients (Rho) between the survey method’s Shannon’s scores were: Crops and weeds: 0.00 (P = 0.96), Tall grass and herb: 0.23 (P = 0.19), Fertile grassland: 0.31 (P = 0.00), Infertile grassland: 0.12 (P = 0.10), Lowland wooded: 0.13 (P = 0.16), Upland wooded: 0.39 (P = 0.04), Moorland grass mosaic: 0.28 (P = 0.02), and Heath and bog: 0.38 (P = 0.00). Four of the AVCs (Fertile grassland, Upland wooded, Moorland grass mosaics, Heath and bog) demonstrate significant (P ≤ 0.05) association between the two survey methods however, the Rho coefficients do not exceed 0.39, indicating ‘weak’ or ‘moderate’ associations. Diversity metrics built on community-level relative abundances did not appear to correlate well between the two survey methods.

Similarly, the richness of genera detected by molecular and 1 m2 surveys for each site was assessed (Figure 2). Average richness scores per site were lower in the molecular survey, except for ‘Crops and weeds’ habitats. The total genera recorded by each method within each AVC were calculated and compared, and the genera that occurred in both survey types were counted as co-recorded. Unique genera values by AVC were, for molecular and 1m^2^ surveys: Crops and weeds: 69 and 38 (27 co-recorded), Tall grass and herb: 42 and 42 (25 co-recorded), Fertile grassland: 70 and 38 (31 co-recorded), Infertile grassland: 88 and 68 (49 co-recorded), Lowland wooded: 28 and 28 (15 co-recorded), Upland wooded: 29 and 39 (17 co-recorded), Moorland grass mosaics: 41 and 42 (24 co-recorded), Heath and bog: 23 and 29 (14 co-recorded). Within the AVC ‘Crops and weeds’, the molecular method records higher richness; in the former, this is likely due to both the residue of previous crops and the detection of ephemeral weeds. The total genera recorded by the molecular method in ‘Fertile-’ and ‘Infertile grassland’ was also higher than traditional methods, where surveying dense grass swords may result in lower richness being recorded. Across all samples and all AVCs, the total richness of genera was 158 and 151 for molecular and 1m^2^ surveys, respectively, with 110 genera co-recorded.

### Co-recorded genus abundance correlations between survey methods

To assess the ability to predict the amount of flowering plant cover from molecular abundance data, comparisons were made against the 1 m^2^ plant survey data. For this assessment, only genera that were co-recorded within each AVC by both survey types were compared using Spearman’s rank correlation of the total percentage abundances of each genus. The coefficients of abundance correlations were: Crops and weeds: 0.51 (P = 0.01), Tall grass and herb: 0.37 (P = 0.07), Fertile grassland: 0.66 (P = 0.00), Infertile grassland: 0.75 (P = 0.00), Lowland wooded: 0.39 (P = 0.15), Upland wooded: 0.75 (P = 0.00), Moorland grass mosaic: 0.58 (P = 0.00), Heath and bog: 0.19 (P = 0.52). Five of the AVCs (Crops and weeds, Fertile grassland, Infertile grassland, Upland wooded, Moorland grass mosaics) demonstrate significant (P ≤ 0.05) association between the survey methods, with Rho coefficients > 0.50 the abundance associations were ‘moderate’ to ‘good’, indicating that where taxa are co-recorded abundances are generally well correlated (Figure 3).

**Figure 3.**
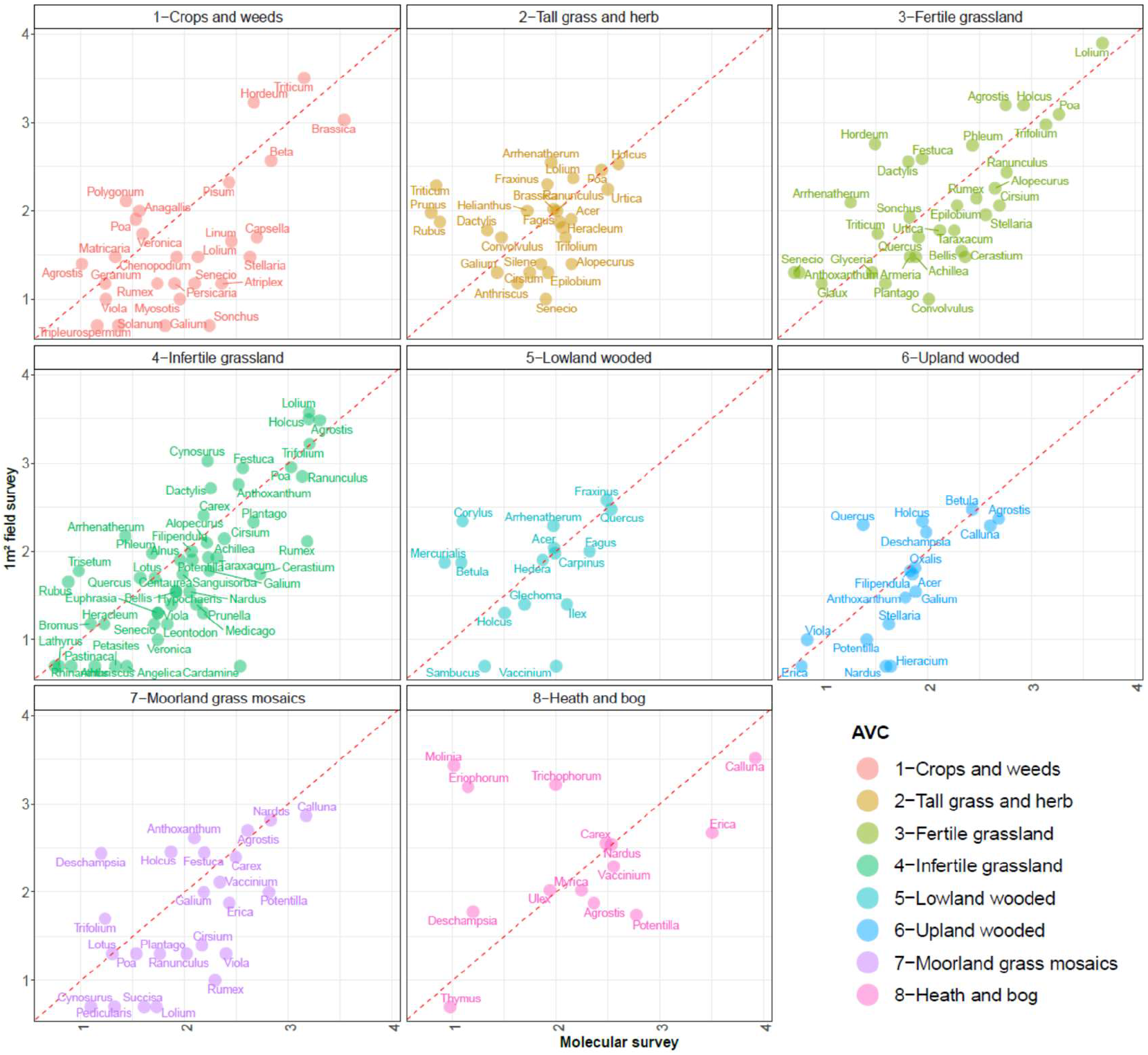
Scatter plots of all co-recorded flowering plant genera across all sites for each AVC habitat. Axis are log^10^ cumulative percentage of genera’s abundance, X-axis molecular, Y-Axis 1m^2^ plant cover, red dotted 1:1 line.

### Genus level indicators of Aggregate Vegetation Classification

From the genus-level abundance data, we assessed the commonality of indicator taxa between survey methods to determine whether molecular data could provide relevant taxonomic discriminators of AVC habitats. Indicator genera with significance (P < 0.05) were determined for each AVC from each survey type. Table 1 lists the significant indicators per AVC ranked by the indicator values (indval). Genera names in bold are those recorded as significant indicators by both survey methods. As per the earlier diversity and richness metrics, the molecular data records more indicators of ‘Crops and weeds’ than the 1m^2^ plant survey data, where the common weeds of agricultural or disturbed land are also recorded as indicators (genera Aegilops, Capsella, Vicia and Atriplex). The molecular data also recorded the same indicator taxa for ‘Fertile grasslands’ and more indicator taxa for ‘Infertile grasslands’ possibly due to difficulty surveying dense grass swards. In the remaining AVC classes the 1 m^2^ plant survey data produced more indicators of vegetation type.

**Table 1.** Aggregate vegetative class indicator genera determined by Molecular and 1m^2^ plant cover surveys. The indicators are listed per AVC habitat and ranked by indicator value (indval) with significance value (p-val). Genera names given in bold with an asterix are those recorded as significant indicators by both survey methods.

| AVC | Molecular survey |  |  |  | 1m<sup>2</sup> field survey |  |  |  |
| --- | --- | --- | --- | --- | --- | --- | --- | --- |
|  | Rank | Genera | indval | p-val | Rank | Genera | indval | p-val |
| 1-Crops and weeds | 1 | Brassica<sup>a</sup> | 0.381 | 0.001 | 1 | Triticum<sup>a</sup> | 0.263 | 0.001 |
|  | 2 | Triticum<sup>a</sup> | 0.249 | 0.001 | 2 | Hordeum<sup>a</sup> | 0.114 | 0.003 |
|  | 3 | Aegilops | 0.101 | 0.001 | 3 | Brassica<sup>a</sup> | 0.076 | 0.008 |
|  | 4 | Capsella | 0.079 | 0.003 | 4 | Beta<sup>a</sup> | 0.041 | 0.016 |
|  | 5 | Hordeum<sup>a</sup> | 0.072 | 0.003 |  |  |  |  |
|  | 6 | Vicia | 0.056 | 0.02 |  |  |  |  |
|  | 7 | Beta<sup>a</sup> | 0.047 | 0.03 |  |  |  |  |
|  | 8 | Atriplex | 0.033 | 0.044 |  |  |  |  |
| 2-Tall grass and herb | 1 | Urtica<sup>a</sup> | 0.115 | 0.002 | 1 | Urtica<sup>a</sup> | 0.172 | 0.001 |
|  |  |  |  |  | 2 | Arrhenatherum | 0.083 | 0.015 |
|  |  |  |  |  | 3 | Heracleum | 0.044 | 0.009 |
|  |  |  |  |  | 4 | Anthriscus | 0.043 | 0.012 |
|  |  |  |  |  | 5 | Brassica | 0.039 | 0.043 |
| 3-Fertile grassland | 1 | Lolium<sup>a</sup> | 0.344 | 0.001 | 1 | Poa<sup>a</sup> | 0.132 | 0.003 |
|  | 2 | Poa<sup>a</sup> | 0.139 | 0.001 | 2 | Phleum<sup>a</sup> | 0.116 | 0.001 |
|  | 3 | Phleum<sup>a</sup> | 0.049 | 0.029 | 3 | Lolium<sup>a</sup> | 0.503 | 0.001 |
| 4-Infertile grassland | 1 | Agrostis<sup>a</sup> | 0.110 | 0.006 | 1 | Trifolium<sup>a</sup> | 0.269 | 0.001 |
|  | 2 | Rumex | 0.100 | 0.002 | 2 | Agrostis<sup>a</sup> | 0.257 | 0.001 |
|  | 3 | Ranunculus<sup>a</sup> | 0.094 | 0.006 | 3 | Holcus<sup>a</sup> | 0.249 | 0.001 |
|  | 4 | Trifolium<sup>a</sup> | 0.094 | 0.021 | 4 | Cynosurus<sup>a</sup> | 0.245 | 0.001 |
|  | 5 | Holcus<sup>a</sup> | 0.081 | 0.014 | 5 | Ranunculus<sup>a</sup> | 0.128 | 0.001 |
|  | 6 | Cerastium | 0.053 | 0.042 | 6 | Dactylis | 0.079 | 0.017 |
|  | 7 | Plantago<sup>a</sup> | 0.050 | 0.028 | 7 | Plantago<sup>a</sup> | 0.057 | 0.017 |
|  | 8 | Cynosurus<sup>a</sup> | 0.047 | 0.023 |  |  |  |  |
| 5-Lowland wooded | 1 | Quercus<sup>a</sup> | 0.217 | 0.001 | 1 | Rubus | 0.253 | 0.001 |
|  | 2 | Fraxinus<sup>a</sup> | 0.203 | 0.001 | 2 | Fraxinus<sup>a</sup> | 0.227 | 0.001 |
|  | 3 | Ilex<sup>a</sup> | 0.143 | 0.001 | 3 | Hedera<sup>a</sup> | 0.190 | 0.001 |
|  | 4 | Hedera<sup>a</sup> | 0.140 | 0.001 | 4 | Quercus<sup>a</sup> | 0.166 | 0.001 |
|  | 5 | Fagus | 0.139 | 0.001 | 5 | Mercurialis<sup>a</sup> | 0.126 | 0.001 |
|  | 6 | Arrhenatherum | 0.063 | 0.009 | 6 | Crataegus | 0.124 | 0.001 |
|  | 7 | Carpinus<sup>a</sup> | 0.048 | 0.033 | 7 | Corylus<sup>a</sup> | 0.123 | 0.001 |
|  | 8 | Mercurialis<sup>a</sup> | 0.048 | 0.029 | 8 | Glechoma<sup>a</sup> | 0.095 | 0.001 |
|  | 9 | Sambucus<sup>a</sup> | 0.048 | 0.026 | 9 | Ilex<sup>a</sup> | 0.095 | 0.003 |
|  | 10 | Glechoma<sup>a</sup> | 0.047 | 0.026 | 10 | Viola | 0.088 | 0.003 |
|  | 11 | Corylus<sup>a</sup> | 0.045 | 0.009 | 11 | Acer<sup>a</sup> | 0.058 | 0.008 |
|  | 12 | Acer<sup>a</sup> | 0.041 | 0.026 | 12 | Elytrigia | 0.054 | 0.018 |
|  | 13 | Nicotiana | 0.039 | 0.03 | 13 | Sambucus<sup>a</sup> | 0.048 | 0.024 |
|  | 14 | Tripleurospermum | 0.037 | 0.046 | 14 | Ballota | 0.048 | 0.017 |
|  |  |  |  |  | 15 | Ulmus | 0.048 | 0.023 |
|  |  |  |  |  | 16 | Rosa | 0.048 | 0.033 |
|  |  |  |  |  | 17 | Carpinus<sup>a</sup> | 0.048 | 0.026 |
|  |  |  |  |  | 18 | Circaea | 0.032 | 0.042 |
| 6-Upland wooded | 1 | Betula<sup>a</sup> | 0.181 | 0.001 | 1 | Deschampsia | 0.101 | 0.006 |
|  | 2 | Ficaria | 0.050 | 0.005 | 2 | Oxalis | 0.086 | 0.001 |
|  |  |  |  |  | 3 | Betula<sup>a</sup> | 0.081 | 0.004 |
|  |  |  |  |  | 4 | Ulex | 0.043 | 0.014 |
| 7-Moorland grass mosaics | 1 | Potentilla<sup>a</sup> | 0.194 | 0.001 | 1 | Nardus<sup>a</sup> | 0.343 | 0.001 |
|  | 2 | Nardus<sup>a</sup> | 0.176 | 0.001 | 2 | Anthoxanthum | 0.300 | 0.001 |
|  | 3 | Carex<sup>a</sup> | 0.090 | 0.007 | 3 | Juncus | 0.256 | 0.001 |
|  | 4 | Viola | 0.073 | 0.011 | 4 | Carex<sup>a</sup> | 0.147 | 0.001 |
|  |  |  |  |  | 5 | Potentilla<sup>a</sup> | 0.134 | 0.001 |
|  |  |  |  |  | 6 | Festuca | 0.095 | 0.015 |
|  |  |  |  |  | 7 | Galium | 0.056 | 0.022 |
|  |  |  |  |  | 8 | Vaccinium | 0.052 | 0.026 |
|  |  |  |  |  | 9 | Sagina | 0.029 | 0.034 |
| 8-Heath and bog | 1 | Calluna<sup>a</sup> | 0.450 | 0.001 | 1 | Eriophorum | 0.477 | 0.001 |
|  | 2 | Erica<sup>a</sup> | 0.381 | 0.001 | 2 | Trichophorum | 0.434 | 0.001 |
|  |  |  |  |  | 3 | Calluna<sup>a</sup> | 0.429 | 0.001 |
|  |  |  |  |  | 4 | Molinia | 0.365 | 0.001 |
|  |  |  |  |  | 5 | Erica<sup>a</sup> | 0.223 | 0.001 |
|  |  |  |  |  | 6 | Narthecium | 0.129 | 0.001 |

### Inventory of recorded genera by survey method

A summary of genus-level abundance data is shown in Figure 4. It displays a complete inventory of the flowering plant genera recorded by each survey method, along with summary statistics for each AVC, including the number of sample sites, the total richness of genera recorded and of which the number that are co-recorded, Spearman’s correlation scores of abundances and average AVC Shannon’s diversity scores. The abundance scores from each survey method (log^10^ of percentage abundance) provided the basis for the heat map.

**Figure 4.**
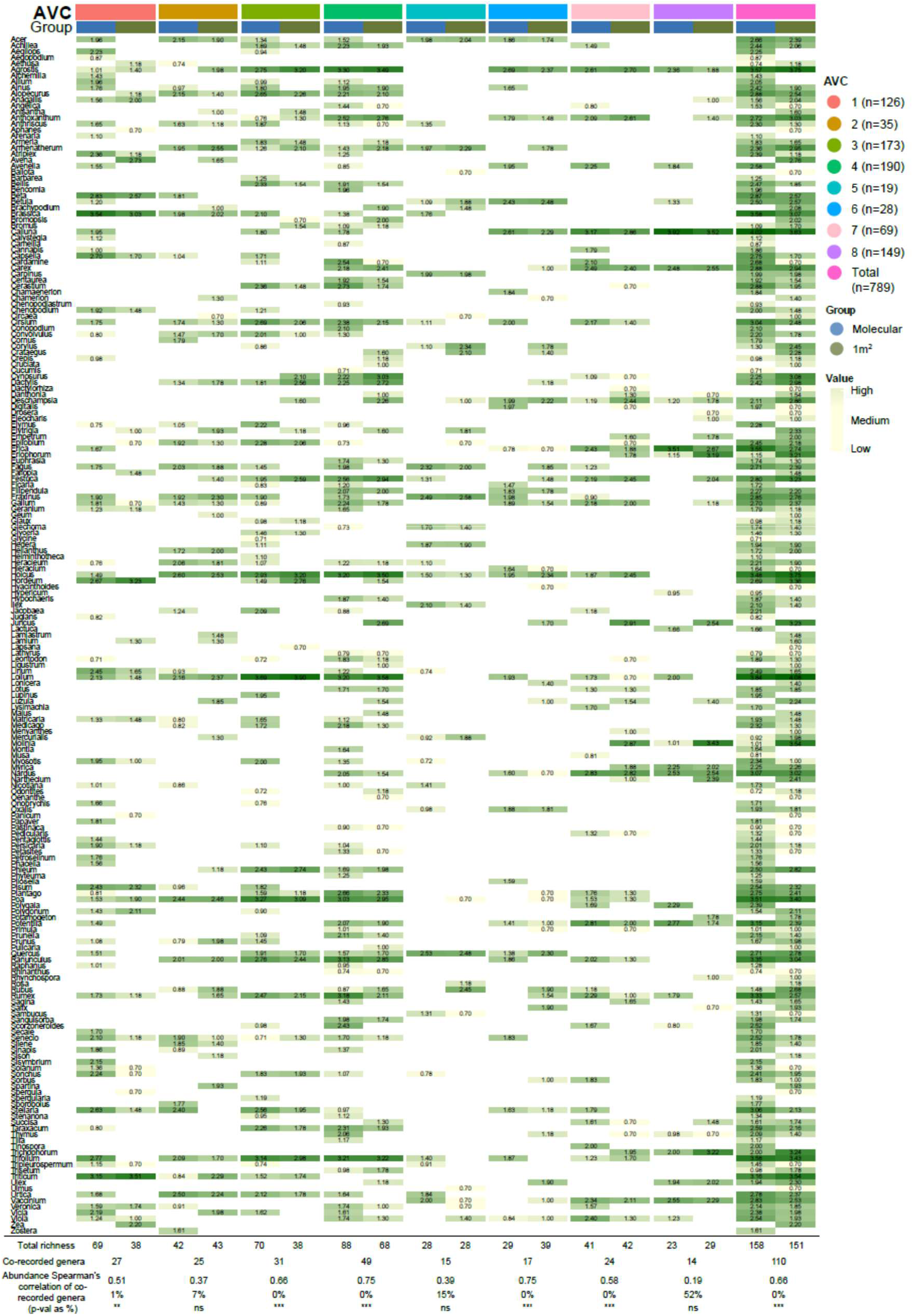
Flowering plant genera inventory and heat map showing occurrence and abundance of all genera recorded by Molecular and 1m^2^ plant cover surveys, with summary statistics for each AVC, including the number of sample sites, the total richness of genera recorded and of which the number that are co-recorded, Spearman’s correlation scores of abundances and average AVC Shannon’s diversity scores. The abundance scores from each survey method (log^10^ of percentage abundance) provide the basis for the heat-map. AVC classes: 1 ‘Crops and weeds’, 2 ‘Tall grass and herb’, 3 ‘Fertile grassland’, 4 ‘Infertile grassland’, 5 ‘Lowland wooded’, 6 ‘Upland wooded’, 7 ‘Moorland grass mosaics’, and 8 ‘Heath and bog’.

### Aggregate Vegetation Classification prediction through machine learning

Abundance data, at the most sensitive taxa level of species, were used to train and validate XGBoost models for the prediction of each sample’s AVC classification from either eDNA or 1 m2 plant cover survey-derived data. Across all AVCs cross validation confusion matrix accuracy for molecular eDNA output was 0.65 compared to 0.75 for 1 m^2^ plant cover survey. With the exception of woodlands and ‘Tall grass and herb’, the predictive power, measured by sensitivity (how many of the actual positive cases we were able to predict correctly), specificity (how many of the correctly predicted cases actually turned out to be positive) and accuracy (how often the classifier correctly predicts) was found to be broadly comparable between the survey types (Figure 5).

**Figure 5.**
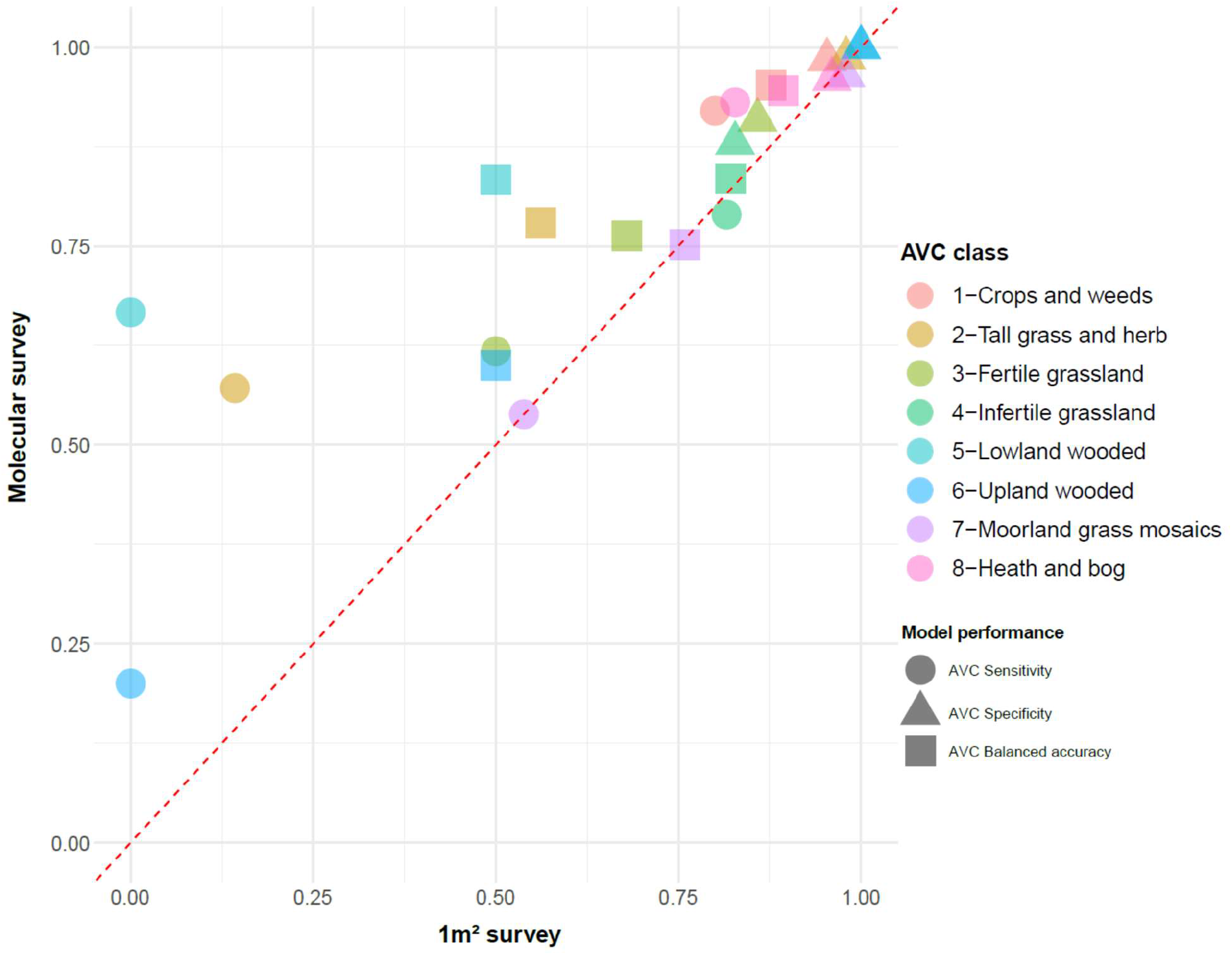
Scatter plot showing degree of predictability of 1m^2^ plant cover (X-axis) and molecular (Y-axis) surveys in AVC assignment. Point colour = AVC classification; point shape = model’s predictive sensitivity (circle), specificity (triangle), and accuracy (square).

## Discussion

Temporal large-scale survey programs encompassing the collection of ecological variables, such as soil state, land use, and plant cover, improve our understanding of the significance, causes, and consequences of large-scale ecological change^3^.

The relative ease of sample collection for eDNA analysis has huge potential for large-scale surveys and citizen science schemes, and could arguably assist in expanding a survey’s range, allowing a more comprehensive inventory to be taken^16^. However, there are many caveats to consider when undertaking eDNA surveys: DNA is a stable biomolecule^17^, and differential persistence and degradation of DNA may present biases in the data. Soil properties can influence the sequestration and persistence of biological molecules. Indeed, the stability of DNA in soil has been shown to depend on moisture, temperature, management, exposure to UV, clay particle type and size and pH^18,19^. Similarly, post-sampling soil storage can have an effect on DNA therein for similar reasons, and the consistency of the storage method should be applied to all samples within a study^20^. Options to this end would include freezing, drying, freeze-drying or the use of proprietary storage buffers such as Zymo research DNA/RNA shield. Other biases, which are particularly pertinent for studies looking for the presence of rare or invasive species, are the relative ease of transmission of eDNA from the site of initial deposition, through a transportive phase (hydrology or disturbance), to a place of persistence^21^, ultimately leading to a study site’s contamination^22^. It is also important to note that plant surveys are not immune from inaccuracies.

Plant characteristics (such as small size, rarity, and morphological confusion), along with environmental factors (such as topography and inclemency) and observer variability, contribute and combine to introduce significant variation and bias, which should ideally be quantified as a quality indicator^23-25^.

DNA has been shown to persist in temperate soils at very detectable levels for >60 years^17,26^, and it is this persistence that may help explain the high diversity measures for AVC-1 ‘crops and weeds’, where the legacy DNA of previous crops and weeds enhances the diversity of plants described by the eDNA method. This persistence raises some questions regarding the time scale needed to detect vegetation change, that is, the sensitivity to land use change. Recently, Foucher et al.^26^ examined eDNA from soil samples in plots for which the crop rotation history was documented and found that the last grown crop formed the dominant taxa in the eDNA inventory, alongside variable detection of past crops up to 8 years with relic-eDNA from historic grape-vines also present. Detection of legacy eDNA may also be advantageous, as it negates the effect of ephemeral genera, those that are small and easily overlooked, or those that are particularly difficult to identify. Indeed, habitat reconstruction through examination of ancient eDNA can facilitate our understanding of lost landscapes and migrations^22,27^.

On average, the eDNA molecular approach used in this study recorded lower genus level diversity scores and richness scores. Similarly, the eDNA approach produced fewer vegetation type indicators. However, within the ‘Crops and weeds’ vegetation type eDNA recorded greater richness and more indicator taxa. It is worth noting that within ‘Crops and weeds’ no correlation was observed for diversity, indicating that eDNA methods are likely recording taxa through the presence of relic-eDNA. This detection of old DNA could be crucial for assessing cropping histories or land use change, but also highlights the potential to record biases in soil eDNA methods due to the variables associated with eDNA persistence within the soil matrix. The molecular data performed well in producing indicator taxa for ‘Fertile’ and ‘Infertile’ -grassland habitats, highlighting more indicators for ‘Infertile grasslands’, possibly due to increased difficulty of surveying thoroughly through dense grass swards.

Using the co-recorded genera of each AVC, we examined the potential of using amplicon-derived relative abundance data to estimate plant cover, a tall ask given the possibility of ploidy, copy number variation, and PCR amplification biases (length and GC content)^28^. Given these concerns, we were surprised that for the majority of vegetation types, the relationship between molecular-derived relative abundance and plant survey cover was moderate to good, demonstrating that it is possible to infer the plant cover of many genera from eDNA abundance data on the proviso that the habitat and genera are appropriate.

To assess the ability to predict AVC from species-level data i.e. the most sensitive level, we employed a machine learning approach to each survey method and found through cross validation that, overall, the traditional survey approach performed better than the molecular eDNA data. This result was mainly driven by ‘-wooded’ and ‘Tall grass and herb’ vegetation types and the inclusion of leaf litter and bare ground in the traditional survey’s ground cover scores, the inclusion of these pseudo taxa providing discriminatory power lacking from the molecular data. Indeed, where vegetation classifications incorporate estimates of the coverage of bare ground, leaf litter or surface rock, molecular data will lose resolution through their omission.

In concordance with the findings reported by Vasar et al.^29^, our molecular results are *broadly* in agreement with those of traditional surveys; however, the level of agreement varied by habitat. In this study, the reduced sensitivity of the molecular data is perhaps unsurprising given the extremely small sample size, what is perhaps surprising is the ability of the eDNA from 0.25 grams of soil to describe the immediate above ground vegetation with some accuracy. Increasing the sample size, pooling from a larger area, or increasing the sample number^30^ would increase the resolving power of the eDNA method. However, there are undesirable cost (increase) and throughput (decrease) implications that need to be weighed carefully. Once collected and stored, soil eDNA can be utilised to examine a range of taxonomic and ecological profiles^31^. The relative ease of sample collection and processing, and the large amount of genetic information held within soil-eDNA makes surveys based on high-throughput eDNA methods very appealing.

The results presented here demonstrate that flowering plant communities and habitats can be described with a degree of accuracy using molecular methods based on small soil-eDNA samples, and that in the absence of field surveys, a molecular approach can provide useful and valuable ecological insights. Collectively, these findings are critical for understanding the efficiencies and disparities between traditional and molecular biomonitoring methodologies. Our national-scale analysis provides insights that highlight the utility of these methods, both traditional and molecular, for large-scale ecological biodiversity measurements.

## Acknowledgements and funding

This project was funded by UKCEH under the ASSIST programme (NERC Reference:NE/N018125/1).

## References

1 Belaire, J. A. et al. Fine-scale monitoring and mapping of biodiversity and ecosystem services reveals multiple synergies and few tradeoffs in urban green space management. Science of The Total Environment 849, 157801 (2022). 10.1016/j.scitotenv.2022.157801

2 Franklin, J., Serra-Diaz, J. M., Syphard, A. D. & Regan, H. M. Global change and terrestrial plant community dynamics. Proceedings of the National Academy of Sciences 113, 3725–3734 (2016). 10.1073/pnas.1519911113

3 Wood, C. M. et al. Long-term vegetation monitoring in Great Britain – the Countryside Survey 1978–2007 and beyond. Earth Syst. Sci. Data 9, 445–459 (2017). 10.5194/essd-9-445-2017

4 Donaldson, L., Wilson, R. J. & Maclean, I. M. D. Old concepts, new challenges: adapting landscape-scale conservation to the twenty-first century. Biodiversity and Conservation 26, 527–552 (2017). 10.1007/s10531-016-1257-9

5 Hiiesalu, I. et al. Plant species richness belowground: higher richness and new patterns revealed by next-generation sequencing. Molecular Ecology 21, 2004–2016 (2012). 10.1111/j.1365-294X.2011.05390.x

6 Cruzan, M. B. et al. Small unmanned aerial vehicles (micro-UAVs, drones) in plant ecology. Applications in Plant Sciences 4, 1600041 (2016). 10.3732/apps.1600041

7 Sharma, R. C., Hara, K. & Hirayama, H. A Machine Learning and Cross-Validation Approach for the Discrimination of Vegetation Physiognomic Types Using Satellite Based Multispectral and Multitemporal Data. Scientifica 2017, 9806479 (2017). 10.1155/2017/9806479

8 Ruppert, K. M., Kline, R. J. & Rahman, M. S. Past, present, and future perspectives of environmental DNA (eDNA) metabarcoding: A systematic review in methods, monitoring, and applications of global eDNA. Global Ecology and Conservation 17, e00547 (2019). 10.1016/j.gecco.2019.e00547

9 Thomsen, P. F. & Willerslev, E. Environmental DNA – An emerging tool in conservation for monitoring past and present biodiversity. Biological Conservation 183, 4–18 (2015). 10.1016/j.biocon.2014.11.019

10 Fahner, N. A., Shokralla, S., Baird, D. J. & Hajibabaei, M. Large-Scale Monitoring of Plants through Environmental DNA Metabarcoding of Soil: Recovery, Resolution, and Annotation of Four DNA Markers. PLOS ONE 11, e0157505 (2016). 10.1371/journal.pone.0157505

11 Carey, P. D. W. S.; Chamberlain, P. M.; Cooper, A.; Emmett, B. A.; Maskell, L. C.; McCann, T.; Murphy, J.; Norton, L. R.; Reynolds, B.; Scott, W. A.; Simpson, I. C.; Smart, S. M.; Ullyett, J. M.. Countryside Survey: UK Results from 2007. 105 (2008).

12 Maskell, L. C., Norton, L.R., Smart, S.M., Scott, R., Carey, P.D., Murphy, J., Chamberlain, P.M., Wood, C.M., Bunce, R.G.H. and Barr, C.J.. CS Technical Report No.2/07 Vegetation Plots Handbook (UK Centre for Ecology and Hydrology, 2008).

13 BA Emmett, Z. F., PM Chamberlain, R Giffiths, R Pickup, J Poskitt, B Reynolds, E Rowe, D Spurgeon, P Rowland, J Wilson, CM Wood CS Technical Report No.3/07 Soils Manual (UK Centre for Ecology and Hydrology, 2008).

14 Bunce, R. G. H. B. C. J.; Gillespie, M. K.; Howard, D. C.; Scott, W. A.; Smart, S. M.; Van de Poll, H. M.; Watkins, J. W.. Vegetation of the British countryside - the Countryside Vegetation System. ECOFACT Volume 1. Vol. 1 224 (UKCEH, 1999).

15 Chen, T. & Guestrin, C. in Proceedings of the 22nd ACM SIGKDD International Conference on Knowledge Discovery and Data Mining 785–794 (Association for Computing Machinery, San Francisco, California, USA, 2016).

16 Biggs, J. et al. Using eDNA to develop a national citizen science-based monitoring programme for the great crested newt (Triturus cristatus). Biological Conservation 183, 19–28 (2015). 10.1016/j.biocon.2014.11.029

17 Yoccoz, N. G. et al. DNA from soil mirrors plant taxonomic and growth form diversity. Molecular Ecology 21, 3647–3655 (2012). 10.1111/j.1365-294X.2012.05545.x

18 Cai, P., Huang, Q., Zhang, X. & Chen, H. Adsorption of DNA on clay minerals and various colloidal particles from an Alfisol. Soil Biology and Biochemistry 38, 471–476 (2006). 10.1016/j.soilbio.2005.05.019

19 Strickler, K. M., Fremier, A. K. & Goldberg, C. S. Quantifying effects of UV-B, temperature, and pH on eDNA degradation in aquatic microcosms. Biological Conservation 183, 85–92 (2015). 10.1016/j.biocon.2014.11.038

20 Clasen, L. A., Detheridge, A. P., Scullion, J. & Griffith, G. W. Soil stabilisation for DNA metabarcoding of plants and fungi. Implications for sampling at remote locations or via third-parties. Metabarcoding and Metagenomics 4, e58365 (2020).

21 Harrison, J. B., Sunday, J. M. & Rogers, S. M. Predicting the fate of eDNA in the environment and implications for studying biodiversity. Proceedings of the Royal Society B: Biological Sciences 286, 20191409 (2019). 10.1098/rspb.2019.1409

22 Pedersen, M. W. et al. Ancient and modern environmental DNA. Philosophical Transactions of the Royal Society B: Biological Sciences 370, 20130383 (2015). 10.1098/rstb.2013.0383

23 Morrison, L. W. Observer error in vegetation surveys: a review. Journal of Plant Ecology 9, 367–379 (2016). 10.1093/jpe/rtv077

24 Ullerud, H. A., Bryn, A., Halvorsen, R. & Hemsing, L. Ø. Consistency in land-cover mapping: Influence of field workers, spatial scale and classification system. Applied Vegetation Science 21, 278–288 (2018). 10.1111/avsc.12368

25 Verheyen, K. et al. Observer and relocation errors matter in resurveys of historical vegetation plots. Journal of Vegetation Science 29, 812–823 (2018). 10.1111/jvs.12673

26 Foucher, A. et al. Persistence of environmental DNA in cultivated soils: implication of this memory effect for reconstructing the dynamics of land use and cover changes. Scientific Reports 10, 10502 (2020). 10.1038/s41598-020-67452-1

27 Haile, J. et al. Ancient DNA Chronology within Sediment Deposits: Are Paleobiological Reconstructions Possible and Is DNA Leaching a Factor? Molecular Biology and Evolution 24, 982–989 (2007). 10.1093/molbev/msm016

28 Álvarez, I. & Wendel, J. F. Ribosomal ITS sequences and plant phylogenetic inference. Molecular Phylogenetics and Evolution 29, 417–434 (2003). 10.1016/S1055-7903(03)00208-2

29 Vasar, M. et al. Metabarcoding of soil environmental DNA to estimate plant diversity globally. Frontiers in Plant Science 14 (2023). 10.3389/fpls.2023.1106617

30 Alsos, I. G. et al. Plant DNA metabarcoding of lake sediments: How does it represent the contemporary vegetation. PLOS ONE 13, e0195403 (2018). 10.1371/journal.pone.0195403

31 Deiner, K. et al. Environmental DNA metabarcoding: Transforming how we survey animal and plant communities. Molecular Ecology 26, 5872–5895 (2017). 10.1111/mec.14350

